# Soil community assembly varies across body sizes in a tropical forest

**DOI:** 10.1101/154278

**Authors:** Lucie Zinger, Pierre Taberlet, Heidy Schimann, Aurélie Bonin, Frédéric Boyer, Marta De Barba, Philippe Gaucher, Ludovic Gielly, Charline Giguet-Covex, Amaia Iribar, Maxime Réjou-Méchain, Gilles Rayé, Delphine Rioux, Vincent Schilling, Blaise Tymen, Jérôme Viers, Cyril Zouiten, Wilfried Thuiller, Eric Coissac, Jérôme Chave

**Author notes:** Ecole Normale Supérieure, PSL Research University, CNRS, Inserm, Institut de Biologie de l’Ecole Normale Supérieure (IBENS), Plant and Diatom Genomics team, F-75005 Paris, France. Univ. Savoie Mont Blanc, CNRS, Ministère de la Culture et de la Communication, EDYTEM, F-73000 Chambéry, France.

## Abstract

The relative influence of deterministic niche-based (i.e. abiotic conditions, biotic interactions) and stochastic-distance dependent neutral processes (i.e. demography, dispersal) in shaping communities has been extensively studied for various organisms, but is far less explored jointly across the tree of life, in particular in soil environments. Here, using a thorough DNA-based census of the whole soil biota in a large tropical forest plot, we show that soil aluminium, topography, and plant species identity are all important drivers of soil richness and community composition. Body size emerges as an important feature of the comparative ecology of the different taxa at the studied spatial scale, with microorganisms being more importantly controlled by environmental factors, while soil mesofauna rather display random spatial distribution. We infer that niche-based processes contribute differently to community assembly across trophic levels due to spatial scaling. Body size could hence help better quantifying important properties of multitrophic assemblages.

## INTRODUCTION

Over a century of documenting patterns of diversity has shown that niche-related, demographic, dispersal and evolutionary processes are all important determinants of ecological communities^1-3^. However, their relative contribution across spatial scales and organisms remains less well known^2,4^, especially when it comes to the soil biota.

Soils are structurally complex environments^5^. They are often described as the “poor man's tropical rainforest”^6^ due to the large and elusive diversity of organisms they harbour^7^. Amplicon-based DNA analysis of environmental samples, or metabarcoding^8^ has recently enabled unravelling novel macroecological patterns for soil fauna, nematodes, bacteria, fungi, protists, and archaea across biomes and habitats^7,9-11^. These patterns often co-vary with environmental conditions such as pH and nutrient quality/availability. They also depend on plant cover due to trophic and mutualistic/pathogenic interactions^12-14^, but this relationship appears context-dependent^15-17^. Also, dispersal limitation or stochastic processes have been unveiled in e.g. fungi or meiofauna^10,18,19^, although other studies reported the predominance of niche determinism^11,20^.

Much of the difficulty in drawing general conclusions about soil community assembly lies in that studies often focus on single taxonomic groups (but see^17,20-22^) or consider different spatial scales. In addition, soil organisms harbour large differences in life-history traits; in particular their body size, whose spans six orders of magnitude (0.1 μm to 10 cm^7^). This property has important consequences for community assembly because organisms of different body size are ruled by contrasting metabolic and demographic processes^23,24^. Dispersal of microorganisms is mediated by external agents and is assumed to occur across large spatial distances. This, together with their large population size and short generation time, would limit local extinction and ecological drift, resulting in a homogeneous global microbial species pool^20,25^. However, microorganisms are also more responsive to small-scale changes in resources, subtle variations that may not be perceived by larger organisms^24,26,27^. Hence, microbial communities are often reported to be dominated by niche-based processes^25^. In contrast, larger organisms (e.g. mesofauna) are thought to be limited in their ability to disperse and have longer reproduction times. They are hence more prone to ecological drift and could display spatial aggregation patterns irrespective of environmental conditions (^28^ and references within). This has enormous consequences for the spatial scaling of soil biodiversity and the major biogeochemical cycles and ecosystem services they sustain^7,12,14^. However, existing knowledge on the scaling of soil communities across body sizes is based entirely on meta-analyses^14,26,27^ and is therefore indirect. This hampers our ability to predict the future of this important biological component^29,30^.

In order to assess the processes governing soil community assemblages, we combined an extensive characterization of abiotic and plant cover conditions with a comprehensive survey of soil biodiversity using DNA metabarcoding in a large tropical forest plot, where samples were collected every ten meters. We sought to (i) determine which factor, both abiotic (e.g. soil chemistry) and biotic (plant diversity, identity), influence soil communities composition, (ii) evaluate the relative importance of niche-based versus neutral processes in shaping these communities and (iii) determine how these effects depend on taxon body size. We predicted that in communities of small-bodied organisms, niche based-processes would be more important relative to larger organisms, which would display spatial aggregation patterns that are consistent with neutrality.

## RESULTS

We found a total of 2,502 archaeal, 19,101 bacterial, and 11,470 eukaryotic OTUs within the 12-ha plot (Table 1). Many of these OTUs were rare (relative abundance < 0.1%) and accounted for 36 ± 10 % (SD values here and after) of archaea OTUs, 86 ± 2 % of bacterial OTUs, and 80 ± 6 % of eukaryotic OTUs. Bacterial OTUs identified at the phylum level (88% of OTUs, 94% of reads) corresponded mainly to Actinobacteria, Alphaproteobacteria, and Acidobacteria. Identified eukaryotic OTUs (51% of OTUs, 70% of reads) belonged to fungi (mainly Agaricomycetes, Glomeromycetes and Orbiliomycetes; Supplementary Table 1), arthropods (mainly termites, mites and springtails) and annelids (Oligochaeta). Plants represented only 2% of all OTUs (10 % of reads). For Archaea, the identified OTUs (3% of OTUs, 64% of reads) corresponded to Nitrososphaeria and Methanomicrobia. Figure 2 illustrates the spatial variation in local diversity for the focal taxonomic groups. The pairwise correlation of these spatial patterns, both in terms of OTU diversity and compositional turnover, was moderate to low (|Pearson’s *r*| < 0.7) and showed low or no correlation with plant ones (Supplementary Table 2-3).

**Table 1.** Soil biota characteristics in the 12 ha plot. Relative abundances were calculated for each marker separately and correspond to the % of reads of each clade. Richness values correspond to the number of OTUs (mean ± SD for values per sample). Diversity values correspond to the effective number of OTUs (exponential Shannon diversity index; mean ± SD for values per sample). Occupancy corresponds to the averaged proportion of samples where an OTU is detected (± SD). NA: not applicable.

| Clade | Total rel. abundance (%) | Total Richness (per sample) | Total Diversity (per sample) | OTU occupancy (%) | Propagule size (m) |
| --- | --- | --- | --- | --- | --- |
| Archaea | NA | 2,502 (43 ± 30) | 30 (9 ± 8) | 1.707 ± 6.197 | 0.5e<sup>-6</sup> |
| Bacteria | NA | 19,101 (1,391 ± 232) | 885 (453 ± 102) | 7.281 ± 16.379 | NA |
| *Acidobacteria* | 19.81 | 1,601 (231 ± 46) | 132 (80 ± 32) | 14.42 ± 22.686 | 0.5e<sup>-6</sup> |
| *Actinobacteria* | 20.578 | 1,696 (167 ± 37) | 86 (54 ± 13) | 9.869 ± 20.329 | 5e<sup>-6</sup> |
| *Bacteroidetes* | 1.023 | 767 (31 ± 12) | 97 (22 ± 8) | 4.088 ± 11.389 | 0.5e<sup>-6</sup> |
| *Chloroflexi* | 5.22 | 2,147 (136 ± 48) | 268 (73 ± 29) | 6.341 ± 12.358 | 1e-6 |
| *Firmicutes* | 2.658 | 873 (19 ± 6) | 4 (3 ± 2) | 2.147 ± 8.048 | 1e-6 |
| *Alphaproteobacteria* | 19.865 | 3,234 (303 ± 57) | 242 (129 ± 25) | 9.373 ± 19.381 | 0.5e<sup>-6</sup> |
| *Betaproteobacteria* | 3.469 | 482 (45 ± 13) | 24 (14 ± 7) | 9.392 ± 19.023 | 5e<sup>-6</sup> |
| *Deltaproteobacteria* | 4.364 | 1,357 (75 ± 31) | 73 (34 ± 15) | 5.513 ± 14.073 | 0.5e<sup>-6</sup> |
| *Gammaproteobacteria* | 8.589 | 888 (72 ± 11) | 30 (21 ± 5) | 8.109 ± 19.47 | 5e<sup>-6</sup> |
| *Verrucomicrobia* | 4.989 | 353 (51 ± 12) | 33 (22 ± 5) | 14.357 ± 24.53 | 0.5e<sup>-6</sup> |
| Eukaryota | NA | 11,470 (475 ± 137) | 228 (51 ± 23) | 4.138 ± 11.113 | NA |
| *Ascomycota* | 3.486 | 317 (25 ± 9) | 22 (8 ± 3) | 7.813 ± 15.365 | 100e<sup>-6</sup> |
| *Basidiomycota* | 16.941 | 387 (31 ± 8) | 44 (8 ± 4) | 7.911 ± 16.184 | 10e<sup>-6</sup> |
| *Glomeromycota* | 1.078 | 38 (7 ± 2) | 8 (5 ± 1) | 19.431 ± 28.548 | 200e<sup>-6</sup> |
| *Annelida* | 6.623 | 49 (8 ± 3) | 6 (3 ± 1) | 17.182 ± 23.828 | 20e<sup>-3</sup> |
| *Arthropoda* | 17.648 | 1,777 (63 ± 21) | 91 (15 ± 7) | 3.569 ± 9.393 | 10e<sup>-3</sup> |
| *Nematoda* | 0.469 | 359 (9 ± 5) | 35 (6 ± 2) | 2.64 ± 7.257 | 100e<sup>-6</sup> |
| *Platyhelminthes* | 0.538 | 117 (4 ± 3) | 15 (3 ± 1) | 3.361 ± 9.778 | 20e<sup>-3</sup> |
| *Protists* | 1.947 | 1,610 (53 ± 40) | 126 (27 ± 18) | 3.29 ± 8.488 | 100e<sup>-6</sup> |

**Figure 1.**
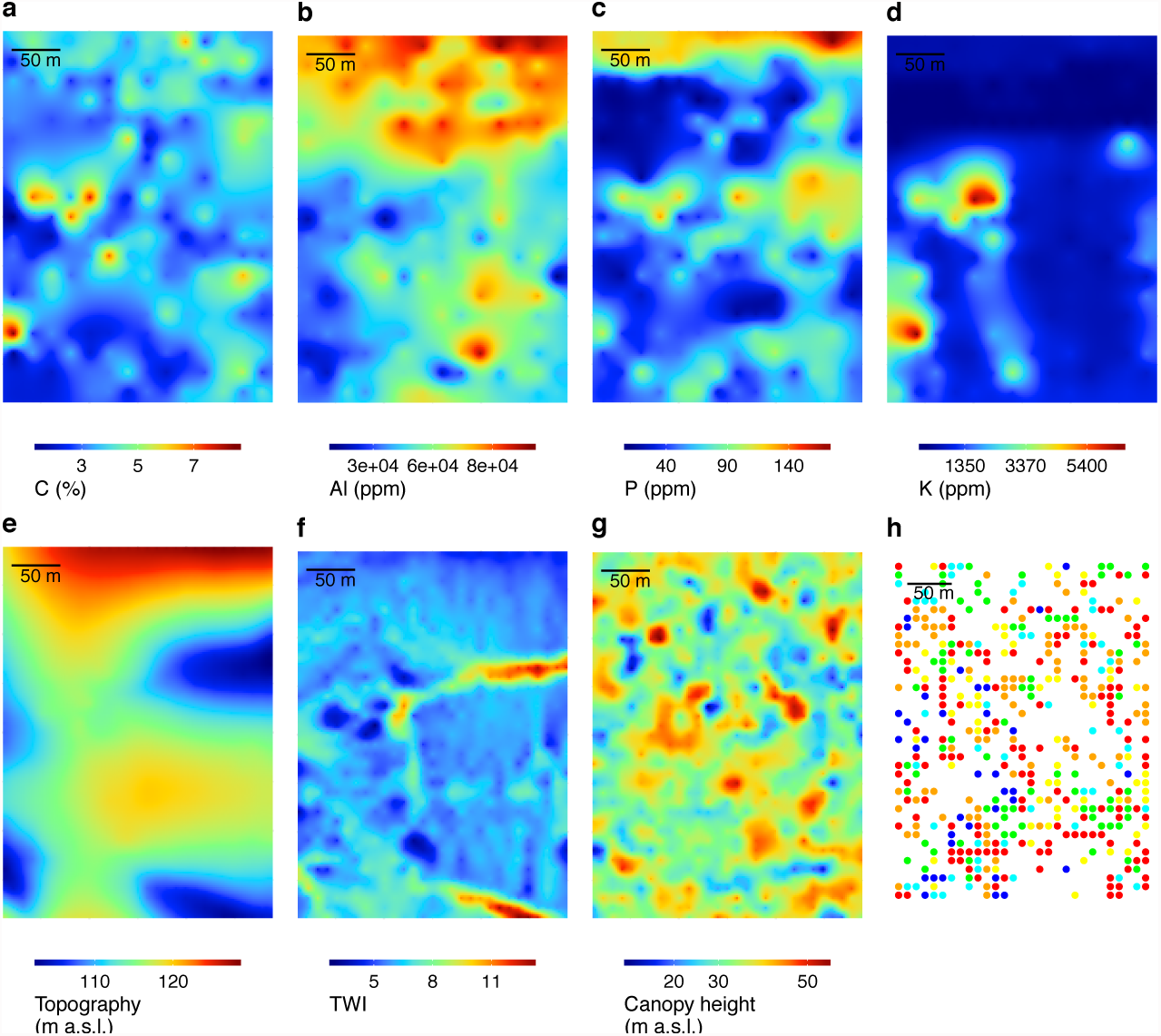
Maps of environmental characteristics in the 12 ha plot. **a-d)** Soil content in Carbon, Aluminium, Phosphorous and Potassium. **e**) site topography, **f**) soil wetness (TWI index, unitless) and **g**) canopy height as inferred from LiDAR data. **h**) Distribution of the most dominant plant genus found in each soil sample as inferred from the plant molecular dataset. Only the six most frequent dominant genera are shown: red: Apocynoideae; orange: Andira; yellow: Ingeae; green: Brosimum; lightblue: Sapotaceae; blue: Drypetes. x/y axes are not shown for representation purposes and range between 10-290 and 10-390 meters respectively.

**Figure 2.**
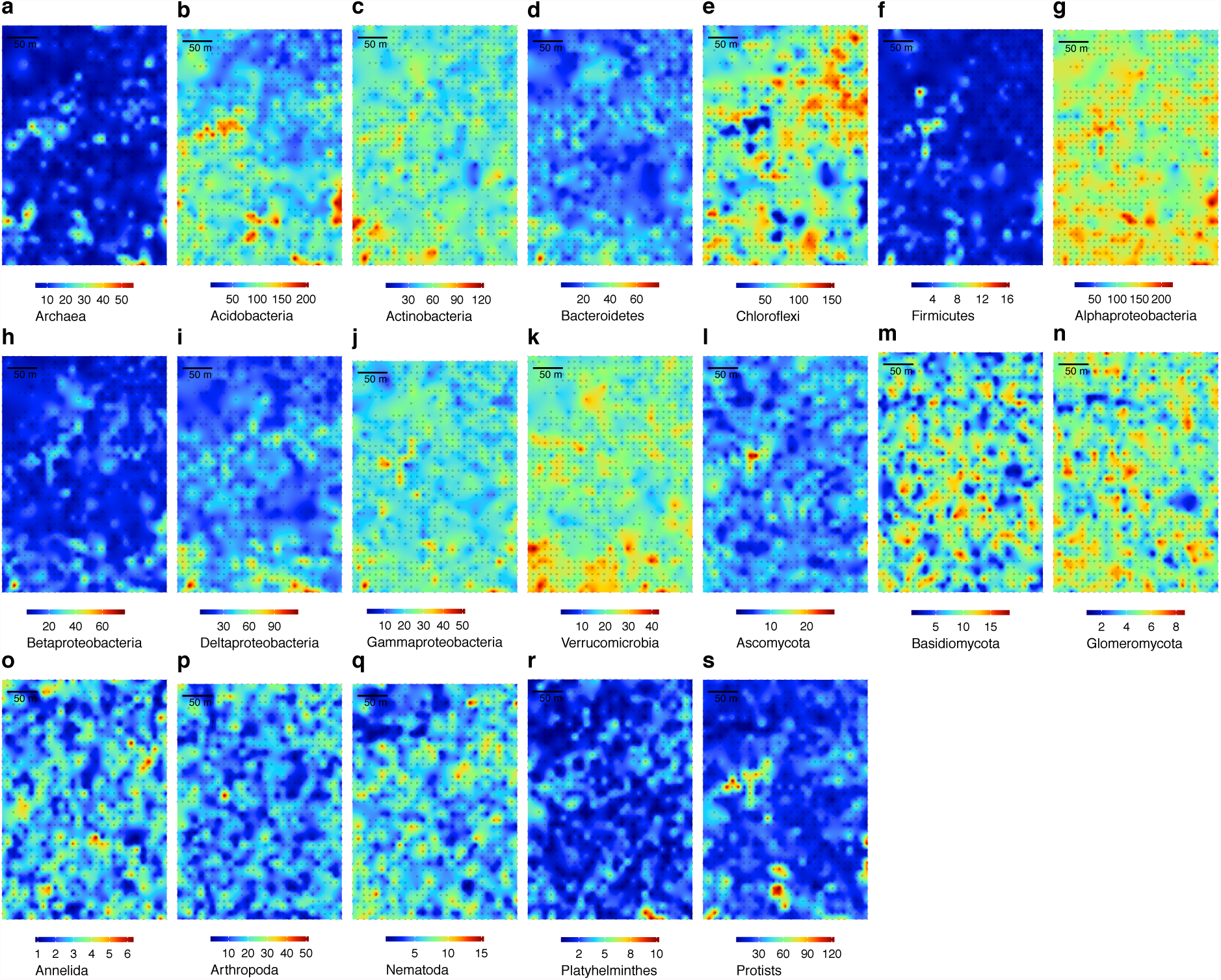
Spatial distribution of OTU diversity per focal clade. The colour scale is expressed as effective numbers of OTUs (exponential Shannon diversity index) for archaea (**a**), bacterial clades (**b-k**) and eukaryotic clades (**l-s**). Maps were obtained by ordinary kriging. Grey dots represent the samples available for each clade.

The full RDA models, i.e. including abiotic, plant and spatial descriptors, explained 3-13% of the variation in OTU composition across the 19 focal taxonomic groups (Fig. 3, Supplementary Table 4). Spatial total effects contributed for 77 ± 9 % of the total explained variation. Half of the spatial variation was explained by spatially structured abiotic conditions (Fig. 1, Supplementary Table 5), in particular, soil aluminium and topographic variables (i.e. elevation, convexity and wetness) or plant characteristics. They correlated negatively with the OTU diversity of most unicellular taxonomic groups for soil aluminium and of several bacterial groups and worms for topographic variables (Supplementary Fig. 1-2). The other half of the spatial variation corresponded to pure spatial effects (41 ± 15% of the total explained variation). Only 7 ± 2% of the total explained variation was due to pure abiotic effects.

**Figure 3.**
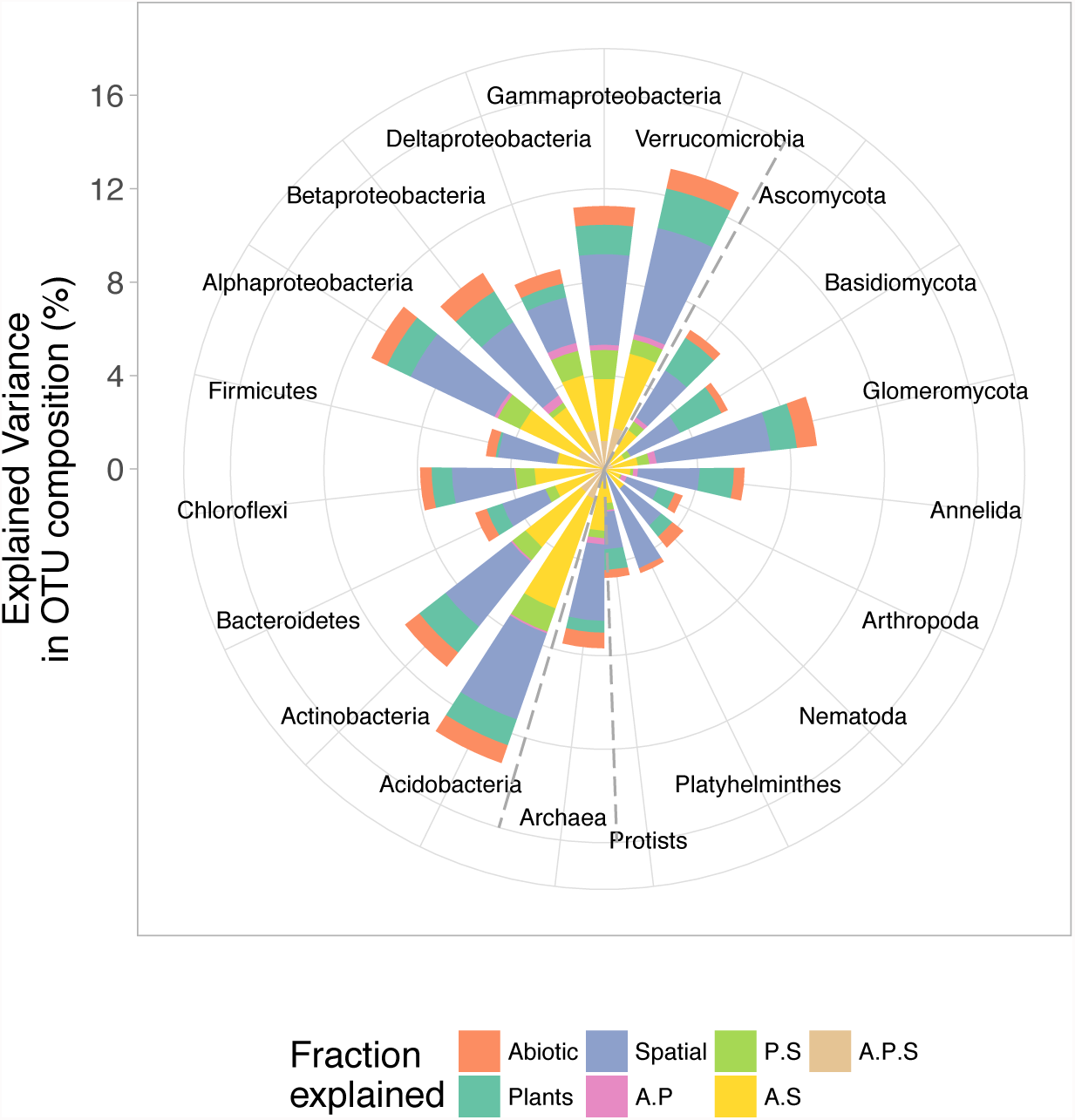
Variation partitioning of OTU composition in each focus taxonomic group. Variations of OTU composition are partitioned into pure (i.e. abiotic (A), plant (P) or spatial (S)) and shared components (A.P, P.S, A.S and A.P.S). The corresponding R^2^_adj_ statistics are reported. See Supplementary Table 4 for corresponding R^2^_adj_ values and their significance (for pure components and full models only, shared components not testable).

Plant total effects also explained a significant fraction of the explained variation (28 ± 11%, Fig. 3) and the plant pure effects were more important than pure abiotic ones in most cases (13 ± 8% of the total explained variation, Supplementary Table 4). The identity of the dominant plant genera in soil samples as detected with the plant DNA marker was the best plant predictors. These variables included 66 different dominant plant genera, 42 of which were unambiguously identified. With OTU diversity instead of OTU composition, 4 to 46% of the variation was explained and the relative importance of plant, abiotic and spatial factors was similar to what observed above (Supplementary Table 6, Supplementary Fig. 3).

Across taxonomic groups, we found a negative relationship between dispersal unit size and the fraction of variation in OTU composition explained by the environmental variables, spatially structured or not (Fig. 4). This relationship held when the full variation or pure abiotic effects were considered alone (Pearson’s *r* = -0.58 and -0.51 respectively, *p* < 0.05). We did not observe such a relationship when performing the analysis on OTU diversity (Supplementary Figure 4).

**Figure 4.**
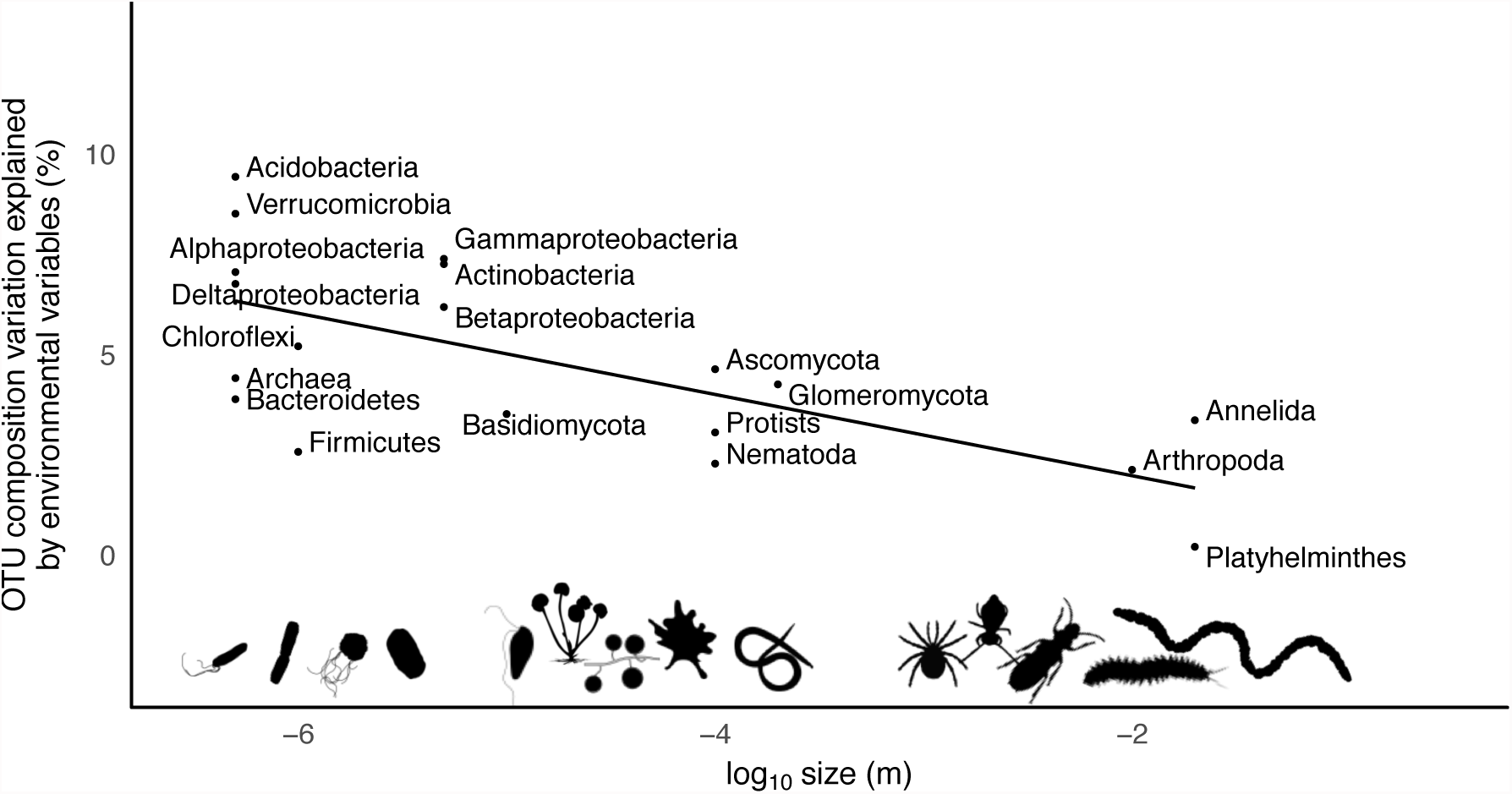
Variation of OTU composition explained by environmental variables according to organism propagule size. The variation (R^2^_adj_ statistics) includes pure plants and abiotic effects as well as their combined effect with spatial variables. The negative relationship between organism propagule size and the variance explained by environmental effects is significant (Pearson’s *r* = -0.67, *p* = 0.002).

## DISCUSSION

Several recent reviews have emphasized the impressive development of publications using high-throughput sequencing to describe the diversity in soils to the extent that it may shift research effort away from fundamental questions in soil ecology and evolutionary biology^13,31^. They emphasize the focus on single taxonomic groups (mainly bacteria/fungi) and the limited consideration for the biotic and abiotic environment at relevant spatial scales are major obstacles to a mechanistic understanding of the processes regulating soil diversity and functioning. This study is an attempt to address these limits, combining assessment of environmental conditions with patterns of soil diversity across the tree of life at relatively reduced spatial scales. We provide evidence for spatial scaling rules across soil taxonomic groups and discuss the general implications of our findings for the assembly of soil communities.

In the spirit of classical community ecology, we sought to understand the spatial distribution of taxonomic groups using environmental and geographical variables. Although the spatial patterns of soil communities differed widely amongst clades, they were all related to several abiotic and biotic factors, albeit to a varying magnitude. This suggests that niche processes always contribute to the community assembly of soil taxa at the studied spatial scale, irrespective of the taxon.

We found total aluminium content to be an important predictor of microbial (i.e. prokaryotes, protists) and nematodes communities. In addition, our study plot is an oxisol with a pH ≤ 5, conditions at which the toxic trivalent Al cation (Al^3+^) predominates over the other forms^32^. Al^3+^competes with other nutrients, inhibits transport, and binds to DNA, ATP or cell walls^32^, as do other metallic ions^33^. In addition, aluminium abundance strongly correlated with that of many other metals (e.g. Ti, Cu, Zn, Cd and Pb, Supplementary Table 4). We hence interpret the lower microbial diversity in Al-rich sites by its toxicity or that of co-varying metals, as suggested for soil tropical microbes^16^. Also, high metal concentration immobilizes nutrients that are essential for plant/microbial growth and protects organic matter from biological degradation^34^. This may explain why total N, C and P contents, which were typical of tropical soils (Methods), were poor predictors of soil diversity. These measures probably do not reflect their bioavailability. High metal conditions likely recruit microbial taxa able to accumulate Al or to exudate organic acids and enzymes to immobilize Al and increase P solubilisation^33,34^. Because soil Al correlates negatively with soil pH (see Methods), our results are also consistent with the decrease of microbial diversity as soil pH decreases repeatedly observed^16,20^. Soil metal content may hence provide a mechanistic understanding of the pervasive soil-pH – microbial diversity relationship, and should be more often considered in addition to other classical measurements.

Topographic variables were also important predictors of OTU composition for many taxonomic groups. They co-vary with many soil properties (e.g. water availability, biogeochemical processes) and most likely reflect meso-habitat heterogeneity. As such, they are often used to predict tropical tree distributions (e.g.^35^) and could be useful to model soil communities. In particular, topographic variables are closely related to soil moisture, and were the most important predictors of OTU diversity of flat/earthworms, known for their sensitivity to this parameter^36,37^, and of some bacterial groups, such as Deltaproteobacteria, an anaerobic clade highly dependent on water availability^38^. This observation generalizes previous findings for soil bacteria^39^ or animals^40^ in tropical environments. We also retrieved high arthropod:earthworm reads abundance ratios consistent with biomass of these groups reported for the dry season^36^, a time where the topographic control in soil humidity is highest and where earthworms migrate deeper into the soil. Accordingly, the arthropod:earthworm reads abundance ratios was lower in a pilot experiment conducted during the wet season. Seasonal dynamics of soil communities should have important implications for biogeochemical cycles, and may be directly impacted by the forecasted increased frequency of Amazonian droughts^41^.

Our environmental parameters further included plant species distribution inferred from DNA metabarcoding. Within our samples, we expected that plant DNA presence relates directly to the quantity/quality of litter or roots exudates^26^, which is poorly controlled by plant diversity/community turnover *per se*^42^. In agreement with this expectation, we found that the locally dominant plant genera influenced the community composition of most focal taxonomic groups (previously termed the “tree identity effect”^7,17^). Neotropical trees species indeed greatly differ in the quantity, quality and chemistry of their litter^43^ and this may hold true for the chemical composition of their root exudates, their influence on soil physical structure or their involvement in specific mutualistic/trophic interactions. This result hence calls for a more mechanistic interpretation of plant-soil interactions^13,14^.

The pervasive contribution of niche-based processes found here echoes previous meta-analyses^44,45^. However, environmental variables explained little variation, whether alone or combined with spatial variables. Previous studies on various soil taxonomic groups reported soil moisture, cation exchange capacity, nutrient or organic matter availability/quality to co-vary with community composition^11,17,20,37^. Although we have characterized the environmental conditions to the best of our ability, we did not quantify directly these important predictors. These often display strong spatial structures in tropical soils^16,35^ and hence potentially contributed to the pure spatial effects observed here. We may also have neglected parameters acting at fine spatial scales, which, together with the high number of samples considered, most likely contribute to the amount of unexplained variance found for all taxa. For instance, we did not quantify soil microstructure, which is important in explaining the small-scale distribution of microbial organisms and water moisture^26,31^, nor did we quantify the geometry of roots as could be imaged by e.g. neutron tomography^5^. We also did not considered soil organisms interactions. These parameters do exhibit considerable fine-scale spatial heterogeneity and might explain the apparent idiosyncrasy of each sample. We might therefore underestimate the contribution of niche-based processes, but expect this to be consistent across the studied soil taxa.

A major problem in attempts to explain patterns of biodiversity is the difficulty of observing ecological processes at the precise scale at which they manifest themselves^4^. Soils are complex media where ecological processes operate at a hierarchy of scales, and any attempt to interpret them should carefully examine the spatial scale of study and the grain of the sampling unit^5,26,27,31^.

For instance, we found no correlation between the species diversity/composition of plants and that of our focal taxonomic groups, contrary to previous reports^15,46^. We interpret this discrepancy as a problem of size and sampling grain. Previous analyses were based on sampling units of typically 20 to 30 m on a side where several soil samples were pooled and compared with aboveground floristic surveys. At this scale, local plant community composition averages over confounding environmental conditions. Our sampling unit was a local soil core of ca. 15 g from which we inferred both local soil richness and plants. Thus, we were able to relate the co-presence of microbial cells and plant cells at the centimetric scale^13^. This allowed us to observe the “tree identity effect” reported above. Recently, Barberan et al. (2015)^16^ reported that the strength of plant-microbial diversity relationship decreased with decreasing sizes of sampling units and vanished at the scale of the sampling point. Our results confirm these findings and extend them to the whole soil biota. Hence, soil biodiversity cannot be inferred from plant floristic patterns at the field scale. As we did not find strong similarities between the spatial patterning of the soil taxonomic groups, we confirm that this conclusion extents to any inference of soil biodiversity patterns from one or few taxa^21^.

Such differences can be explained by strong among-clade differences in body size. Indeed, a striking result from our analysis was that the amount of variation in diversity or composition explained by environmental variables decreased significantly and linearly with increasing propagule size across taxonomic groups.

The smallest body-size category includes bacteria, which was primarily explained by environmental factors, owing to the clear environmental gradient at our site. This is in agreement with the view that niche-based processes dominate bacterial community assembly^25^. Micro-eukaryotes displayed an intermediate amount of explained variation, suggesting niche filtering is less important for these groups than for bacteria. This generalizes previous comparisons of bacteria and fungi community assembly^22^. Whether it is due to longer generation time, lower dispersal ability or broader resources distribution remains to be determined. In fungi, the mycelium can spread across heterogeneous environments and the apparent distribution of fungi may be less related to environmental conditions at the scale of observation. Archaea and certain bacterial groups (i.e. Firmicutes and Bacteroidetes) displayed similar features. These groups are of low abundance in soils or associated with termites^9,16,20,47^, which suggests that they have smaller population sizes in soils or dispersal rates similar to those of macroorganisms, and could be hence more prone to ecological drift.

On the other hand, the OTUs corresponding to large-bodied organisms, i.e. arthropods, annelids and flatworms, had a random spatial distribution. Soil mesofauna assembly may hence be rather determined by neutral processes at the scale of our 12 ha plot and grain studied. However, pure spatial variation, which is usually indicative of dispersal limitation, was less important for this group than for microbes, which contrasts with previous reports^28,44^. We explain this result by the minimal distance between sampling points, 10 m, which was the result of a compromise between spatial resolution and sampling effort. It may have been insufficient to detect spatial aggregates for soil mesofauna, which usually displays spatial aggregation below 10 m^19,26,37^. This roughly corresponds to the horizontal distance earthworms can disperse per year. Alternative sampling strategies should be hence considered for multi-taxa assessments at the studied spatial scale. Still, studies using a finer sampling grain suggest these aggregates to seldom covary with environmental factors and to most likely result from stochastic processes^19,37^. On the opposite end of the range, niche-based community patterns emerge at scales larger than 12-ha where soils properties are highly distinct in tropical trees^35^ or mesofauna^40,48^.

This observation is in line with Levin’s argument on the problem of scales^4^. For a unique studied area and sampling grain, we considered patterns of diversity at a multitude of spatial scales across the studied taxa owing to their body size; with large-scale patterns (i.e. microbes) being more predictable than fine-scale ones (i.e. mesofauna). This result further points to an interesting juncture between two fields of ecological theory that have heretofore been loosely connected, namely the role of body size in explaining the scaling rules of metabolism^23^ and that of environment and dispersal in explaining the assembly of ecological communities^2^. In that light the fact that small-bodied organisms are more influenced by environmental conditions, while large-bodied organisms display neutral assembly is consistent with the main predictions of macroecology^2^ and soil science^26,27^. Our finding of a log-linear relationship between the contribution of niche-based processes in community assembly and propagule size generalizes previous empirical observations in freshwater ecosystems (^28^ and references within). Although other important biological features also likely explain the differences of spatial distribution between soil organisms (e.g. clonal/sexual reproduction, mutualistic/pathogenic interactions, transport that is active, mediated by animals, by water or wind), the body size trait constitutes an operational parameter that cuts across the tree of life. It is indicative of the way organisms perceive – or move across – space^3,24,26,27^ and of their trophic status in its broadest sense^29^. Hence, our result provides empirical evidence of spatial scaling rules across the soil food web.

The recent explosion of DNA-based studies has considerably increased our knowledge on the taxonomic, genetic and functional diversity of soil organisms^7^, but has so far provided limited understanding of the mechanisms shaping soil biodiversity^31^. We do believe that quantifying soil biodiversity using amplicon-based techniques is a useful endeavour. It is increasingly obvious that important properties of multitrophic systems, e.g. species richness, distribution and extinction rates, cannot be easily retrieved from one or a small set of surrogate taxa (this study, ^21,30^). Co-occurrences multitrophic networks can now be reconstructed using such DNA-based approaches^49^, and these approaches hold great promises in assessing soil ecological networks properties at spatial and temporal scales that have been heretofore inconceivable. Our results show that accounting for body size differences helps unravel spatial patterns in such complex communities and, combined with DNA-based approaches, could improve predictive models of soil food webs.

## METHODS

### Study site and sampling

The study site is located at the Nouragues Ecological Research Station, in the lowland rain forest of French Guiana (latitude: 4° 4' 28" N, longitude: 52° 40' 45" W). Rainfall is 2861 mm.y^-1^ (average 1992-2012), with a two-month dry season (< 100 mm.month-1), from late August to early November, and a shorter dry season in March. Our sampling campaign was conducted November 7-20, 2012, towards the end of the dry season, which lasted from early September to late November. Cumulated rainfall during the 60 days preceding the sampling session was 134 mm, with 44 days without rain, and over 90% of the rainfall concentrated in seven days.

We surveyed a 12-ha (300 x 400 m) plot established in 1992. This plot extends on a gentle slope between a ridge and a small creek^50^. The 5,640 trees occurring in the plot (diameter at breast ≥ 10 cm) belong to over 600 species, with the two dominant species accounting each for only 2.3% of individuals^51^. Sand and clay fractions are about 40% each in the soil top 10 cm. The parent material is Caribbean granite. Soil edaphic conditions are typical of tropical oxisols, with an acidic pH (pH = 5.0) and low exchangeable cation content (ECEC = 3.5 cmolc.kg-1). The C:N and N:P ratios are typical of tropical forests^52^ (median=13.4 and 40.5 respectively).

We sampled the plot following a regular grid scheme with a 10-m mesh, excluding bordering points, hence resulting in a total of 1,132 sampling points. At each point, 50-100g soil cores were collected with an auger at ~10 cm depth, excluding the organic horizon. We did so because the organic and mineral horizons harbor different arthropods and microbial communities^53,54^. Lumping together these compartments may hence complicate the interpretation of the spatial distribution of certain soil clades. Consequently, we focused on the surface soil layer, which it is the most biogeochemically active in the mineral horizon^55^. The soil cores were stored and sealed in sterile plastic bags after collection and transported to the field station laboratory. Extracellular DNA was extracted from 15 g of soil per soil core as described previously^56,57^ within 4 hours after sample collection to prevent from microbial growth. DNA was extracted twice for each soil core, and the remaining soil material was dried and stored for analytical chemistry analyses.

### Molecular analyses

Soil biodiversity was surveyed through DNA metabarcoding using four DNA markers, with primers targeting three in hypervariable regions of the *ssu* rRNA gene in Archaea (this study), Bacteria^58^ and Eukaryota^59^ domains respectively (see Supplementary Table 7) and a plastid DNA marker (P6 loop of the trnL intron^60^) to characterize the plant composition at the scale of the sampling point. The universal primers do not present particular amplification biases (see Supplementary Material for a detailed description of primer similarity with priming sites across phyla).

We conducted duplicated PCRs for each marker and each DNA extract, hence representing a total of 18,112 independent PCRs. To discriminate PCR products after sequencing, forward and reverse primers were tagged with a combination of two different 8-nucleotide labels. Each PCR reaction was performed in a total volume of 20 μl and comprised 10 μl of AmpliTaq Gold^®^ Master Mix (Life Technologies, Carlsbad, CA, USA), 0.25 μM of each primer, 3.2μg of BSA (Roche Diagnostic, Basel, Switzerland), and 2 μl DNA template that was 10-fold diluted to reduce PCR inhibition by humic substances. Thermocycling conditions are shown in Appendix S1. All PCR products were then purified using a MinElute^TM^ PCR purification kit (Qiagen, Hilden, Germany). For each marker, PCR products were distributed into 4 different sequencing libraries, which were pooled and loaded on up to 7 Illumina sequencing lanes, depending on the marker and sequencing platform used (Supplementary Table 7), using the paired-end sequencing technology. To control for potential contaminants^61^ and false positive caused by tag-switching events^62^, the sequenced multiplexes comprised extractions/PCR blank controls, as well as unused tag combinations.

### Sequence analyses and curation

The ca. 10^9^ sequencing reads produced were curated using the OBITools package^63^ and R scripts (www.r-project.org).

We first assembled paired-end reads and assigned them to their respective samples and taxonomic groups on the basis of the tags and primer sequences, by allowing 2 and 0 mismatches on primers and tags, respectively. Reads were dereplicated, and low-quality sequences excluded (i.e. shorter than 7, 13, 50 and 100 nt for the plant, eukaryota, archaea and bacterial markers, respectively; containing “Ns”; and singletons). We then clustered the remaining unique sequences into operational taxonomic units (OTUs), as follows. We computed pairwise dissimilarities between sequences (i.e. the number of mismatches, allowed to be 0-3) using the Sumatra algorithm^64^, then we formed OTUs using the Infomap community detection algorithm^65^. We excluded OTUs represented by a single sequence because PCR/sequencing almost always produce at least one error on amplified fragments. The true sequence of an OTU was assumed to be the most abundant one in the cluster.

OTUs were assigned a taxonomic clade with the ecotag program of the OBITools package. Since this algorithm requires full-length barcodes as reference, we constructed a set of reference databases for each marker by running in silico PCRs with the primer pairs used here (Supplementary Table 7). This was done with the ecoPCR program^66^ by using Genbank (release 197; ftp://ftp.ncbi.nlm.nih.gov/genbank) and the MOTHUR-formatted SILVA database (release 119; http://www.mothur.org/wiki/Silva_reference_files) as sequence template. We here only considered references with unambiguous taxonomic annotation at the order level. We also used a reference database of tropical plants occurring at the study site, built with the plant primers, so as to improve plant OTU identification (available in ^56^ and the Dryad Digital Repository, doi:10.5061/dryad.1qt12). At the end of this analysis, we had two taxonomic assignments for each OTU. We gave priority to assignments from SILVA or our local plant database when the similarity between the query and its best match was > 98%. We did so because the SILVA and our local plant databases yield taxonomic annotations that are more reliable than those from Genbank. Otherwise, the taxonomic assignment yielding the highest similarity score was kept. Paired-end reads were assembled, assigned to their respective samples/marker and dereplicated. Low-quality sequences were excluded; the remaining ones were clustered into operational taxonomic units (OTUs) and assigned a taxonomic clade.

We paid particular attention to minimize PCR/sequencing errors, contaminant and false positive sequences as well as potential non-functional PCRs by using several conservative quality criteria. First, we assumed that OTUs peaking in abundance in the negative controls were a contaminant. Any OTU with a best-match similarity in any reference database below < ca. 75% was considered as a chimera or highly degraded sequence. Any OTU that did not fall into the clade targeted by the primer pair was also excluded. Finally, we also curated the dataset from false positives caused by tag-switching events. This phenomenon is suspected to occur during the preparation of the sequencing library and translates into low abundance “contaminant OTUs” in samples coming from other samples that were part of the amplicons multiplex. To remove them, we considered each OTU separately and set to 0 any abundance representing < 0.03% of the total OTU abundance in the entire dataset, similarly to ^62^. This abundance threshold was found to be the one for which most OTUs with unused tag combinations could be removed from the dataset.

Verifying the success of our 18,000 PCRs was not possible. Therefore, we used the PCR replicates to filter out potential non-functional PCR reactions, i.e. containing low amounts of reads or high amounts of contaminants or false positives that could not be filtered out with the process described above. First, for each sample we defined an average community (hereafter centroid) by averaging OTUs counts of the four PCR replicates. Second, we compared the distribution of dissimilarities (as defined by Euclidean distances) of PCR replicates with their respective centroid (hereafter *d_w_*) against the distribution of pairwise dissimilarities between all centroids (hereafter *d_b_*). As we expect *d_w_* < *d_b_*, we defined the intersection of *d_w_* and *d_b_* distribution curves as the dissimilarity level above which a PCR replicate is too distant from its centroid to be reliable. Any PCR above this threshold was excluded from the analysis. This process was repeated iteratively until no more PCR were removed from the dataset. If a sample was represented by only one PCR product, then it was excluded during the iterative process. At the end of this procedure, the remaining PCR replicates were summed for each sample. See Supplementary Table 8 for raw and curated datasets statistics.

Finally, the sequencing depth of each sample was standardized for each marker by randomly resampling a number of reads equal to the first quartile of read number across samples. Although such procedure has been recently questioned^67^, the data loss caused by rarefaction is minimal when the subsampling size well covers the sample diversity, as it is the case here. It had therefore no or weak effects on the dataset characteristics and the retrieve patterns of diversity (Supplementary Fig. 5). Raw and curated sequencing data as well as associated metadata are available on the Dryad Digital Repository under the accession XXX.

### Focus taxonomic groups and body size

Within the Eukaryota and Bacteria domains, we distinguished groups on the basis of their taxonomic affiliation at the phylum level (Table 1). We did so because broadly defined functional traits such as body size and trophic categories are relatively well conserved within phyla^38,68^. We restricted our analysis to the most abundant phyla (i.e. representing ≥ 1% of the total bacterial or eukaryotic OTU diversity). Archaea were analysed as a single group due to imprecise taxonomic assignments.

Body size classes were defined as the size of the dispersal unit (propagule), rather than that of the mature individual: in some taxa, like fungi, mature bodies may extend over large areas through mycelium growth (i.e. the vegetative part of fungi). The intraspecific variability of these mature forms besides vary from 1 to 2 orders of magnitude^69^. This makes it difficult to decide on an effective definition of a ‘body’. In contrast, spore size is more stable and proportional to the reproduction rate and to the fructification size of certain fungal groups^69,70^. This definition also cuts across the domains of life as it corresponds to average cell size for unicellular organisms (i.e. archaea, bacteria and protists) and average body size of adults for the soil meiofauna. For fungi, we used their spore size, as the operational definition of body size^18^. Body sizes were inferred from^71^ for bacterial cells, from^72,73^ for fungal propagules, and from^68^ for the other groups (Table 1).

### Environmental parameters

#### Soil chemistry and airborne lidar

Total content in soil chemical elements was assessed on one soil sample in two (i.e. every 20 meters). Soils were ground and attacked by hydrochloric and nitric acid. Major element concentration was measured by inductively coupled plasma optical-emission spectroscopy (ICP-OES). Carbon and nitrogen concentration was measured by a CHN elemental analyzer (NA 2100 Protein, CE Instruments). All other chemical elements were quantified by ICP-MS. In total, our dataset included 55 elemental concentrations, which were krigged using an exponential variogram model so as to obtain values for all the points of the initial sampling design. This analysis was conducted with the *sp* (<u>http://rspatial.r-forge.r-project.org/</u>) and *gstat* (http://gstat.r-forge.r-project.org/) R packages. Despite soil pH is a common predictor of soil microbes^74^, we did not assess this parameter here. Soil pH is indeed well known to correlate negatively with soil total aluminium, in both temperate^75^ and tropical soils^76^. Because tropical soils are Al-rich, conditions that are extreme for the soil biota^34^, we believe this variable to be more relevant in our study plot than soil pH *per se*, as the latter results from mixed biochemical/chemical pathways and hence provides a poor mechanistic explanation of the processes shaping soil communities. Airborne lidar data were obtained previously^77,78^ and were used here to retrieve a 1-m^2^ digital elevation model (DEM), from which we derived slope, light penetration, and topographic wetness indices (see Supplementary Methods).

#### Plants

We used several plant-related variables to explain soil community assembly. The first corresponded to the canopy closure derived from the lidar data. The other corresponded to plant diversity (Shannon index) and the identity of the three most dominant plant genera in each soil sample as inferred from the plant metabarcoding dataset. Dominant plant genera represented on average 70 ± 15% of the plant reads in each sample, which provides a good description of the local plant community composition. Sampling points for which we did not obtained any plant sequences (330 of the 1,132) were redrawn by a multinomial resampling of reads from neighboring points. This approach is reasonable because tree roots influence is detectable up to 20 m from their corresponding stems in tropical forests^16^.

### Statistical analyses

To disentangle the relative importance of niche-based and neutral processes in the assembly of soil communities, we used variation partitioning and redundancy analysis (RDA^44,79^) on each focus taxonomic group. We used RDA because preliminary analyses indicated a linear relationship between soil diversity/community distribution and explanatory variables. First, we Hellinger-transformed OTU tables to down-weight rare OTUs, as these may be artifactual, as well as to preserves the Euclidean distance among sites^80^. Next, we constructed three parsimonious models by applying a forward selection procedure. This avoids inflating the amount of explained variance and Type I error^81^.

A first model corresponded to RDAs including soil chemistry and lidar-derived data. The second corresponded to RDAs including plant explanatory variables (Fig. 1, Supplementary Table 9). For these two models, explanatory variables were preselected to reduce multicollinearity (|Pearson’s *r*| ≥ 0.7, Supplementary Fig. 6) and normalized with a Box-Cox transformation to meet the normality assumption. The third model was a spatial RDA using spatial eigenvectors as explanatory variables, derived from a Principal Components of Neighbours Matrices (PCNM) approach^82^. These represent spatial structures at different spatial scales that allow modelling the spatial structure of community composition variation. To further reduce inflation of *R^2^* statistics caused by the PCNM analyses^83^, we preselected PCNM eigenvectors prior the forward selection procedure. These corresponded to eigenvectors explaining significantly (p ≤ 0.02) the variability in the biological response, as assessed though partial canonical redundancy analysis (pRDA). This pre-selection was performed for each studied clade independently. Geographic coordinates were also included in the model to account for possible linear trends along the study area^82^.

Abiotic, plant and spatial models were then combined into a single, “full” RDA model, which was subjected to variation partitioning. This analysis decomposes the variance of community composition explained either by abiotic, plant or spatial variables alone as well their combined effects. The contribution of pure environmental vs. spatial effects in the variation of community composition is usually considered to be indicative of the relative importance of niche-base processes vs. dispersal limitation in the community assembly^44,79^, provided that the environmental context is well characterized and that neutral processes are not correlated with the environment. Significance of the total RDA models and pure effects were determined with 1,000 Monte-Carlo permutations. We here only report *R^2^*_adj_ statistics, which are less inflated when the numbers of explanatory variables is high^[84]^. To further identify the environmental parameters that were spatially structured in our plot, each PCNM eigenvector was regressed against the set of plant and abiotic variables using pRDA.

To test an effect of body size on community assembly patterns, we compared it with the amount of variance explained by the full model and each pure effect using a Pearson product-moment correlation test. Finally, we repeated the whole analysis by considering OTU diversity as a response variable. OTU diversity was calculated as the exponential of the Shannon entropy, an effective number of species^85^ that is less sensitive to rare OTUs. All analyses were conducted with the vegan R package (<u>http://vegan.r-forge.r-project.org/</u>).

### Data availability

Raw and curated sequencing data as well as associated metadata and codes are available on the Dryad Digital Repository (XXX provided upon manuscript acceptance).

## Acknowledgements

We are indebted to the staff of the Nouragues Research Field Station (CNRS, French Guiana) and the Nouragues Natural Reserve, for commodities and technical help during the field session. We thank Johan Pansu, Guilhem Sommeria-Klein, Mélanie Roy and Renato Lima for helpful discussions on the manuscript, as well as the Genotoul bioinformatics platform Toulouse Midi-Pyrenees (Bioinfo Genotoul) and Pierre Solbes (EDB-Calc Cluster) for providing computing and storage resources. The work was funded by the METABAR project (ANR-11- BSV7-0020) and has benefitted from "Investissement d'Avenir" grants managed by Agence Nationale de la Recherche (CEBA: ANR-10-LABX-25-01; TULIP: ANR-10-LABX-0041; OSUG@2020: ANR-10-LABX-56; ANAEE-France: ANR-11-INBS-0001). This work is dedicated to the memory of our colleague Serge Aubert.

## Author contributions

L.Z., E.C., P.T. and J.C. conceived the study. L.Z, P.T., H.S., A.B, M.D.B, P.G., L.G., C.G.C, A.I., M.R.M, G.R., E.C and J.C. contributed to the fieldwork and/or to DNA extractions. P.T., A.B., A.I. and D.R. conducted the laboratory work to produce the metabarcoding data. J.V. and C.Z. performed the chemical analyses. B.T. provided the LiDAR data. L.Z. did the bioinformatics and statistical analyses with the help of V.S., F.B., E.C, W.T. and J.C. The manuscript was written by L.Z. and J.C. with input from all co-authors.

